# Parallel erosion of a testis-specific Na^+^/K^+^ ATPase in three mammalian lineages sheds light into the evolution of spermatozoa energetics

**DOI:** 10.1101/2025.04.07.647512

**Authors:** Raul Valente, André Machado, Eva Pericuesta, David García-Parraga, Nádia Artilheiro, Filipe Alves, Isabel Sousa-Pinto, Bernardo Pinto, J. Miguel Cordeiro, Raquel Ruivo, Alfonso Gutiérrez-Adán, L. Filipe C. Castro

## Abstract

Understanding how extant physiological landscapes arise from novel genetic interactions is key to elucidating phenotypic evolution. Sperm cells exemplify a striking case of functional compartmentalization shaped by molecular adjustments, notably regarding energy metabolism. Here, we examine the impact of gene duplication and loss on the evolution of sperm energetics in mammals. Our findings reveal that the acquisition of an exclusive mechanism controlling the sperm plasma membrane Na^+^ gradient, critical for glucose uptake, emerged in the ancestor of mammals through gene duplication, which originated the Na^+^/K^+^ ATPase transporting subunit alpha 4 transporter (*Atp1a4*). Furthermore, we demonstrate that testis-specific expression of *Atp1a4* was acquired after the divergence from monotremes. Notably, we identify three independent pseudogenization events of *Atp1a4*, including in pangolins, the naked mole-rat (*Heterocephalus glaber*) and toothed whales. The recurrent loss of function in *Atp1a4* coincides with the erosion of the testis-specific glycolytic pathway in these lineages. Furthermore, enrichment analysis of striped dolphin (*Stenella coeruleoalba*) and naked mole-rat testis transcriptomes also suggests significant alterations in sperm capacitation processes. Overall, we show that the elaboration of a sodium-dependent glucose uptake wiring was a key innovation in the energetic landscape governing mammalian spermatozoa, with secondary gene loss in three separate lineages pointing to drastic alterations in motility and capacitation processes. Our findings illustrate how metabolic pathways co-shaped by gene duplication and erosion define extant physiological phenotypes.

**Significance:** The sperm cell exhibits a distinct morphology, characterized by exceptional functional and energetic compartmentalization between the midpiece and the flagellum. Here, we demonstrate that the evolution of a sodium-dependent glucose uptake mechanism, driven by gene duplication and giving rise to Atp1a4, was a key innovation promoting efficient glucose usage in mammalian sperm. Additionally, we identify independent losses of Atp1a4 in three lineages—pangolins, naked mole-rats, and toothed whales—coinciding with the reduction of glucose-dependent pathways, such as glycolysis. Our findings highlight how gene expansion and erosion shape metabolic pathways, offering insights into the genetic innovations underlying physiological diversity in reproductive biology.

## Introduction

The diversity and complexity of gene networks are dependent on the intricate dynamics of multiple evolutionary processes (e.g., De Smet and Van de Peer 2012). At the centre stage of gene (re)wirings is the impact of gene duplication and loss events (Albalat and Cañestro 2016; Copley 2020; Sánchez-Serna et al. 2024), contributing to the emergence of novel phenotypic features (Helsen et al. 2019). While gene duplication *offers* new genetic material fostering functional novelty, gene loss can partially or completely eliminate genes or gene hubs, with occasional consequences at the adaptive level (e.g., Huelsmann et al. 2019; Espregueira Themudo et al. 2020; Xu and Guo 2020). The spermatozoon provides a remarkable example of the evolutionary assembly of gene networks. In fact, this fascinating cell type at the very centre of species survival and perpetuation (Puerta Suárez et al. 2018), possesses a unique morphological and functional profile, that is also coordinated at the molecular level (Gu et al. 2019; Brattig-Correia et al. 2024). For instance, sperm cells exhibit a characteristic compartmentalization of energy sources (Miki et al. 2004; Zecchin et al. 2015), with ATP provided by both glycolysis and oxidative phosphorylation (OXPHOS): for short- and long-term usage at the flagellum and midpiece, respectively (Westhoff and Kamp 1997; Storey 2008; Zecchin et al. 2015; Alves et al. 2021). Importantly, such compartmentalization is mirrored by an autonomous sperm cell glycolysis pathway, independent from somatic tissues, partially operated by exclusive gene duplicates (*e.g.,* sperm-specific *Gapdhs*) and alternative splice isoforms (Miki et al. 2004). The exceptionality of this energy network is further strengthened by specific glucose uptake mechanisms. In agreement, glucose transport was recently shown to be critically dependent on the preservation of a proper Na^+^ sperm cell membrane gradient, which is ensured by a Na^+^/K^+^- ATPase subunit alpha 4, *Atp1a4* (Numata et al. 2022a; Numata et al. 2022b). Similarly to the sperm cell glycolysis modules, the mouse *Atp1a4* is specifically expressed in the testis/sperm cells; and, targeted inactivation of the gene generates a sterile male phenotype (Jimenez et al. 2011). How this sophisticated energy network of ATP synthesis, *via* ion-dependent glucose uptake and subsequent glycolysis, evolved in mammalian male reproductive cells is unknown. Here, we show that the sperm-specific Na^+^/K^+^-ATPase, *Atp1a4*, emerged in the ancestor of mammals *via* gene duplication. Our findings suggest that upon duplication this Na^+^/K^+^ ATPase was specifically co-opted into the energy gene network of the Theria mammal ancestor sperm cell. We further show the parallel elimination of spermatozoa glucose uptake *via* gene loss of *Atp1a4*, that took place in the ancestor of toothed whales, the naked mole rat (*Heterocephalus glaber*) and pangolins.

## Results and Discussion

### The Na^+^/K^+^-ATPase, *Atp1a4*, emerged in mammalian ancestry

To investigate the timing of *Atp1a4* origin, we performed a phylogenetic analysis with a broad taxonomic coverage. We included *Atp1a2* gene sequences due to their reported similarity to *Atp1a4* sequences (Shull et al. 1986; Underhill et al. 1999; Keryanov and Gardner 2002). The inferred maximum likelihood tree places the reptile and bird *Atp1a2* genes outside a monophyletic group containing all mammalian *Atp1a2* and *Atp1a4* sequences (Figure 1 - Panel A). The latter were divided into five well-supported groups encompassing Monotremata *Atp1a2*/*Atp1a4*, placental mammal *Atp1a2*, marsupial *Atp1a2*, marsupial *Atp1a4* and placental mammal *Atp1a4* sequences respectively (Figure 1 - Panel A). In addition, comparative synteny analysis indicates that mammalian *Atp1a2* and *Atp1a4* reside in a tandem arrangement (Figure 1). Thus, the appearance of *Atp1a4* coincides with mammalian radiation, as bird and reptile single copy sequence lineages (here represented by the chicken (*Gallus gallus*) and green turtle (*Chelonia mydas*)) outgroup both *Atp1a2*/*Atp1a4* mammalian orthologues (Figure 1 - Panel B). Moreover, Monotremata—an early branching clade of mammals—contains two genes that encode ATPases in the same genomic region, thus suggesting that *Atp1a2/4* gene duplication early mammalian evolution, but sequence divergence happened after the separation of marsupials and placental mammals.

**Figure 1.**
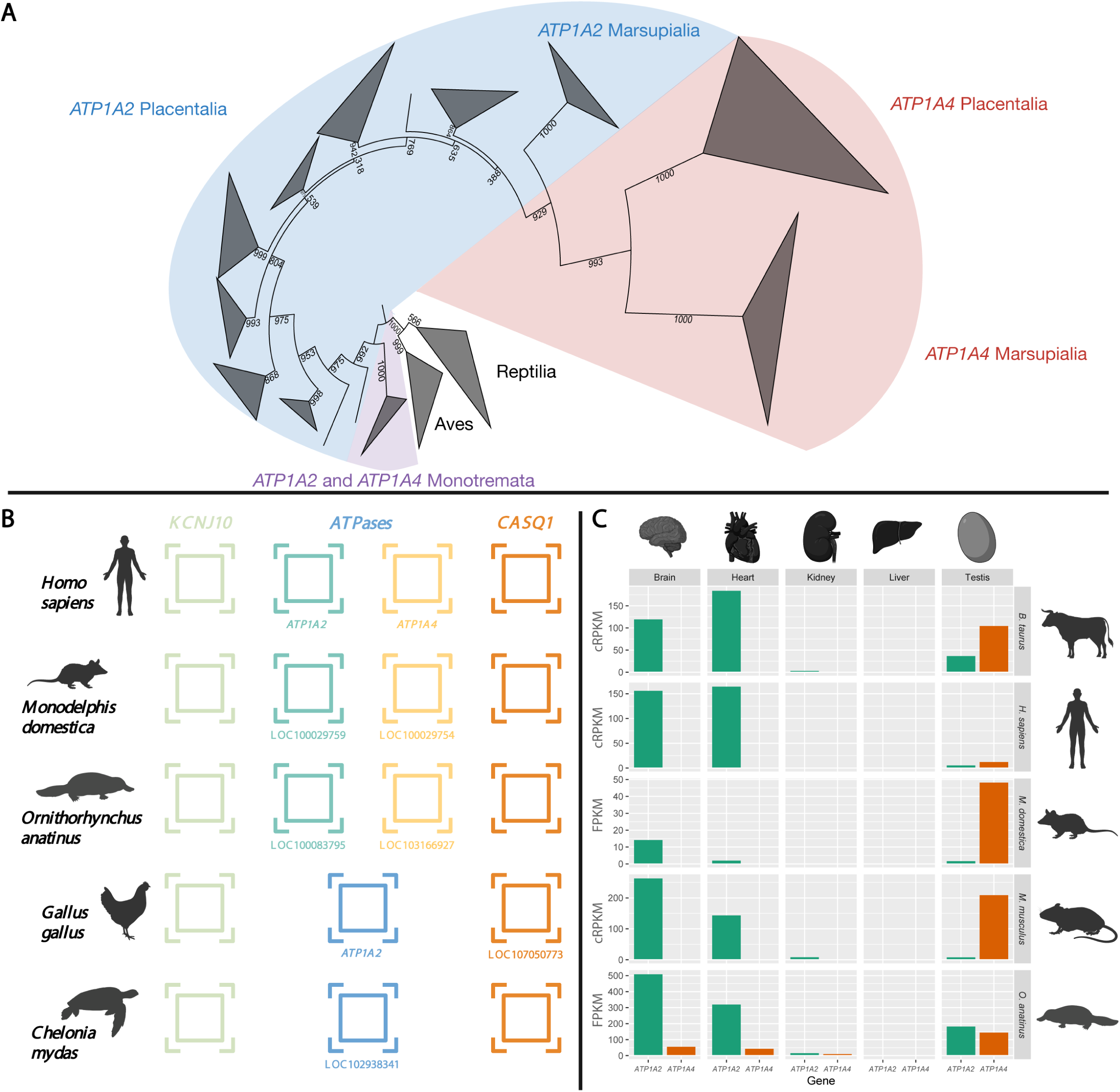
Evolution of *Atp1a2* and *Atp1a4* in the mammalian lineage. A) Maximum Likelihood phylogenetic tree of the *Atp1a4* and *Atp1a2* sequences. Number at nodes represent bootstrap values. B) Synteny of the *Atp1a2/4* locus. C) Gene expression (units in FPKMs or cRPKM) of *Atp1a4* and *Atp1a2* in the platypus, gray short-tailed opossum, human, mouse and bull. Data from the platypus was retrieved from Sequence Read Archive from NCBI. Data from humans, mice, and bulls was retrieved from Vastdb (Tapial et al., 2017).

### *Atp1a4* gene expression is testis-restricted in the Theria lineage

*Atp1a4* was previously established as a testis-specific gene of Na^+^/K^+^-ATPase in mice (Shamraj and Lingrel 1994), with the KO displaying infertility (Syeda et al., 2020). Yet, it is unknown when this tissue specificity emerged and whether it was concurrent with the *Atp1a2/4* gene duplication event. Thus, we explored several multi-tissue RNA-seq datasets, including the platypus (*Ornithorhynchus anatinus*), a species from an early branching mammalian lineage (Monotremata). Our results indicate that the platypus “*Atp1a2/4”* genes are expressed in various tissues. In contrast, *Atp1a4* is predominantly expressed in the testis of humans (*Homo sapiens*), mice (*Mus musculus*), bulls (*Bos taurus*), and the gray short-tailed opossum (*Monodelphis domestica*) (Figure 1 - Panel C). Therefore, our analysis strongly suggests that after the gene duplication of the ancestral *Atp1a2/4* during early mammal evolution, *Atp1a4* expression became specified to testis/sperm cells before the separation of marsupials and placental mammals.

### Signs of sequence erosion of *Atp1a4* at three separate Mammalia lineages

The finding of a functional coupling between a sodium/potassium membrane exchanger, ATP1A4, and a sodium/glucose cotransporter, SGLT1, in mice entails an interesting evolutionary scenario (Numata et al. 2022a; Numata et al. 2022b). For example, it implies the emergence of a specific and efficient mechanism for glucose uptake to support sperm cell motility. Conversely, the removal of glucose as a key energetic substrate for sperm cells, would render a glucose-dedicated sodium/potassium pump useless. Interestingly, sperm glycolysis has been suggested to have undergone erosion in some mammals, including toothed whales (Alves et al. 2021). Thus, we next explored the coding status of *Atp1a4* across 202 mammalian species with publicly available genomes. Genomic regions containing the *Atp1a4 locus* were scanned with Pseudo*Checker*, an automated pipeline that estimates the coding status of a gene by detecting open-reading-frame (ORF) disruptive mutations (Alves et al. 2020). Based on PseudoIndex—a user assisted metric incorporated into Pseudo*Checker* that classifies the coding status of a gene in a discrete scale from 0 (coding) to 5 (pseudogenized)—74 species displayed scores higher than 2, suggesting *Atp1a4* erosion. Within these species, high PseudoIndex values were mostly due to fragmentation of the genomic region (presence of Ns), exon absence, poor alignment identity or scaffold incompleteness in the *Atp1a4* genomic region. Additionally, in some species (e.g., the Bactrian camel (*Camelus ferus)*) further mutational validation did not confirm the disruption of the *Atp1a4* gene.

### Multiple inactivating mutations are found in the *Atp1a4* sequence of Odontoceti Cetacea

Cetaceans are one of the few mammalian groups in which the *Atp1a4* sequence shows robust signs of sequence of erosion. In Mysticeti (baleen whales), four out of eleven species analysed exhibited validated ORF-disrupting mutations. Specifically, minke whales (*Balaenoptera acutorostrata scammoni* and *Balaenoptera bonaerensis*) harboured a conserved one-nucleotide deletion in exon 10, along with other mutations. On the other hand, in the gray whale (*Eschrichtius robustus*) and the fin whale (*Balaenoptera physalus*), the *Atp1a4* coding sequence (CDS) exhibited in-frame premature stop codons and frameshift mutations.

In Odontoceti (toothed whales) the scenario is markedly different, when compared to the sister clade. The mutational landscape suggests a complete impairment of *Atp1a4* coding status across the lineage (Figure 2 - Panel A). Conserved mutational events were identified within Ziphiidae (beaked whales), with a two-nucleotide deletion in exon 19, and in Delphinoidea - a superfamily comprising oceanic dolphins (Delphinidae), true porpoises (Phocoenidae), and the narwhal and beluga (Monodontidae) - where a four-nucleotide insertion in exon 15 was found to be conserved across all members of this superfamily (Figure 2 - Panel A).

**Figure 2.**
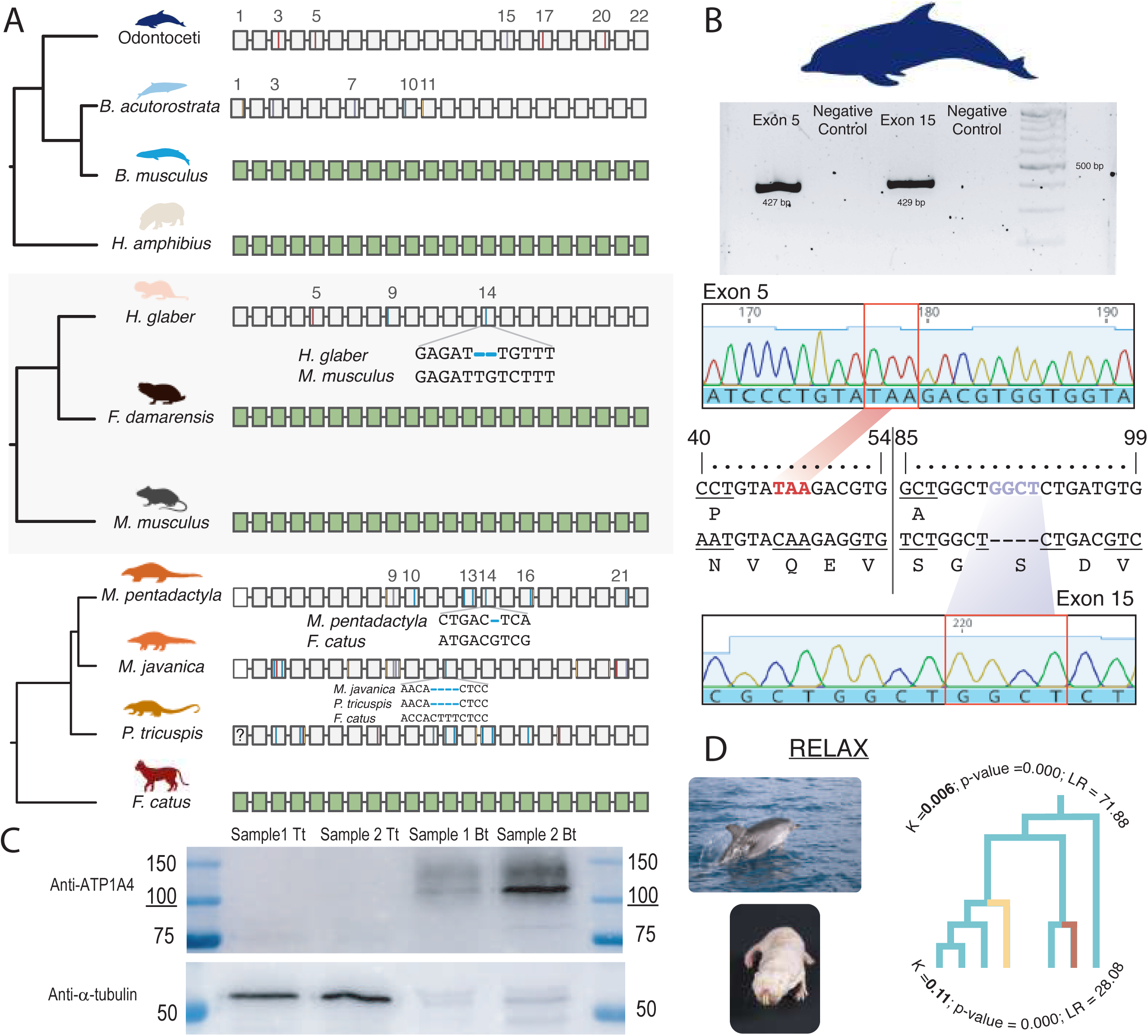
Erosion of *Atp1a4* gene in three mammalian lineages: Cetacea, pangolins and the naked mole rat. A) ORF-disrupting mutational landscape in *Atp1a4* of cetaceans, naked mole-rat and pangolins compared to outgroups (green box – coding exons; insertions in purple, deletions in blue, in-frame premature stop codons in red, and splice site mutations in yellow). B) PCR electrophoresis and electropherogram to validate disruption of *Atp1a4* in *T. truncatus*. C) Western Blot image for ATP1A4 (∼ 100 kDa) and α-tubulin proteins from bottlenose and bull sperm cells. D) RELAX analysis for cetaceans and naked mole-rat. Spotted dolphin (*Stenella frontalis*) photograph taken by Ágatha Gil. Naked mole-rat photograph retrieved from flickr (https://www.flickr.com/).

The observed erosion landscape was further validated for Odontoceti using a stepwise approach. First, with polymerase chain reaction and Sanger sequencing, the conserved four-nucleotide insertion in exon 15, as well as an in-frame premature stop codon in exon 5, were confirmed using genomic DNA obtained from *T. truncatus* skin samples (Figure 2 - Panel B). Concordantly, transcriptomic analysis of striped dolphin (*Stenella coeruleoalba)* testis further supported *Atp1a4* inactivation in Odontoceti, with residual gene expression found for *Atp1a4* (TPM ≅ 0; see below). Lastly, the absence of ATP1A4 protein production was investigated using spermatozoa samples from both common bottlenose dolphin (*T. truncatus*) and bull (*B. taurus*). In agreement, western blot analysis revealed a band corresponding to ATP1A4 in cattle samples (expected size around 100-kDa), whereas in common bottlenose dolphin no protein bands of similar size could be observed (Figure 2 - Panel C).

### The coding gene sequence of *Atp1a4* is impaired in the naked mole-rat but not in Damaraland mole-rat

Besides the non-coding status of *Atp1a4* in Odontoceti cetaceans, we also collected evidence of ORF-disruption in the naked mole-rat (*H. glaber*). More specifically, *Atp1a4* in naked mole-rat exhibited an in-frame premature stop codon in exon 5, a four-nucleotide deletion in exon 9 and a two-nucleotide deletion in exon 14 (Figure 2 - Panel A). In stark contrast with the pattern found in *H. glaber*, Pseudo*Checker* analyses yielded no signs of pseudogenization for Damaraland mole-rat (*Fukomys damarensis*). Yet, further manual annotation of *F. damarensis Atp1a4* revealed the presence of a unique single-nucleotide insertion in the last exon of *Atp1a4*. Nevertheless, in agreement with previous studies suggesting that mutations occurring at the beginning or end of a gene do not necessarily imply gene loss (MacArthur et al. 2012; Sharma et al. 2016), we did not consider this mutation sufficient to confirm the loss of function of *Atp1a4* in Damaraland mole-rat.

### Inactivation of *Atp1a4* across Pholidota species

In addition to cetaceans and naked mole-rat, pangolins also displayed signs of pseudogenization, yielding a PseudoIndex of 5 in all analysed members. Further manual annotation of the ORF-predictions confirmed the poor coding status of *Atp1a4* in these species with multiple disruptive mutations found along the gene (Figure 2 - Panel A). Interestingly, no conserved ORF-disruptive mutations were found among *Manis* sp. (*M. javanica* and *M. pentadactyla*); however, a conserved 4-nucleotide deletion was found in exon 12 in both *Phataginus tricuspis* and *M. javanica*. While Pholidota seem to possess a non-functional *Atp1a4*, all analysed members of its sister clade (Carnivora) retain a coding *Atp1a4* (Figure 2 - Panel A).

### Relaxed selection might have contributed to *Atp1a4* pseudogenization

To elucidate whether the molecular evolution of *Atp1a4* is under relaxed or intensified selection, we performed RELAX tests (Wertheim et al. 2015) using both Cetacea and naked mole-rat as test branches. All other mammals presenting a coding *Atp1a4* were used as reference branch to calculate the relaxation or intensification parameter (k, with k<1 indicating relaxation of selective strength) of *Atp1a4* in each class. The RELAX test suggests a scenario of *Atp1a4* evolution under relaxed selection in both cetaceans and naked mole-rat (k===0.006 [p===0.000] and 0.11 [0.000]) (Figure 2 - Panel D), leading to significant changes in *Atp1a4* amino acidic composition.

### Enrichment analysis in testis highlights shifts in sperm physiological processes in striped dolphin and naked mole-rat

Given the suspected loss of *Atp1a4*, an essential component for sperm cell energy metabolism, we searched for differentially enriched pathways in spermatozoa of naked mole-rat (*H. glaber*) and striped dolphin (*S. coeruleoalba*) in comparison with mouse (*M. musculus*) and bull (*B. taurus*), respectively. Using a comparative transcriptomic approach, we identified the key pathways that were exclusively enriched in the testis of mouse and bull (compared to striped dolphin and naked mole-rat) (Figure 3). Among these were find “*sperm capacitation*” (GO:0048240), “*regulation of cilium movement involved in cell motility*” (GO:0060295), “*regulation of cilium-dependent cell motility*” (GO:1902019), or “*CatSper complex*” (GO:0036128). Interestingly, these GO terms include genes that play significant roles in capacitation: a sequence of sperm physiological and morphological modifications that trigger the ultimate acrosome reaction and allow fertilization (e. g. hyperactivated motility, membrane hyperpolarization and fluidity, cellular content and pH, cytoskeleton remodelling, protein phosphorylation, ROS generation (Rajamanickam et al. 2017)). For instance, the CatSper complex comprises a family of cation channels, functioning through Ca^2+^ signalling, that initiate the tyrosine phosphorylation cascade (Chung et al. 2014), required for sperm motility and fertility in mammals (Ren and Xia 2010). To understand whether these novel network landscapes result from tissue-specific reduction or absence of gene expression or, ultimately, from the loss of genes related to these GO terms requires further investigation. Conversely, GO terms associated with lipid binding and storage (“*lipoprotein particle binding*” (GO:0071813) and “*cholesterol storage*” (GO:0010878)) were enriched in naked mole-rat and striped dolphin in comparison with the references, mouse and bull (Figure 3).

**Figure 3.**
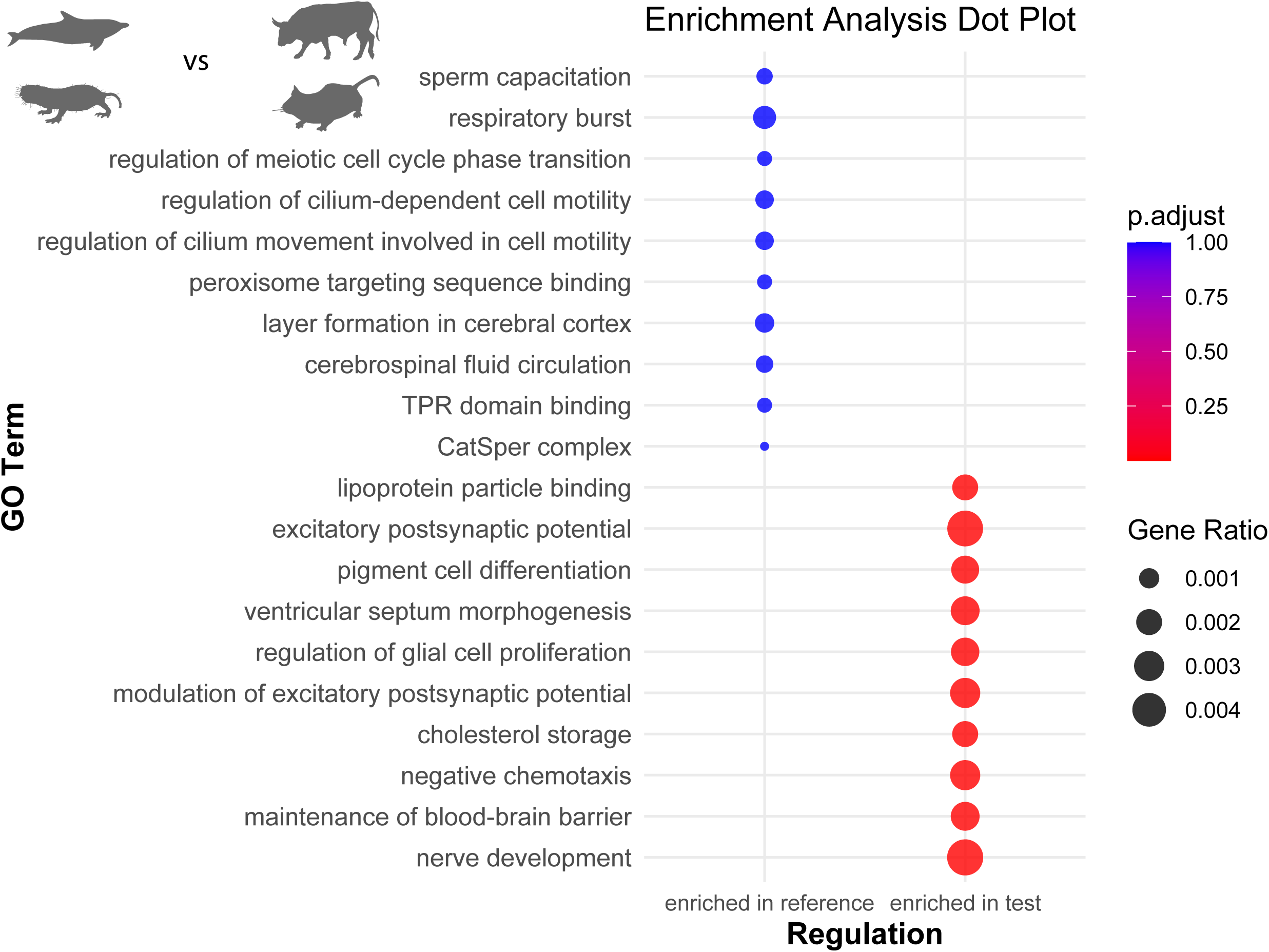
Comparative gene expression in testis across 4 mammal species. Dot plot illustrating the enrichment analysis of testis gene expression in *H. glaber* and *S. coeruleoalba* compared to *M. musculus* and *B. taurus*. The top Gene Ontology (GO) terms are highlighted - 10 terms in each of the following categories: exclusively enriched in *M. musculus* and *B. taurus* (enriched in reference), and exclusively enriched in *H. glaber* and *S. coeruleoalba* (enriched in test).

Overall, our findings suggest a regulatory shift in the molecular machinery associated to sperm physiology. These observations agree with the unique characteristics of naked mole-rat and Cetacea sperm. In naked-mole rat, spermatozoa were found to be structurally simplified and with reduced motility. This degenerate profile was associated with the absence of sperm competition in a eusocial mammal with reproductive patterns mostly restricted to a single female and few breeding males (van der Horst et al. 2011). Additionally, sperm cells of both naked mole-rat and toothed cetaceans were shown to have different energetic requirements, paralleled by the dismantling of glycolysis and an enhanced dependence on fatty acid ß-oxidation (Alves et al. 2021). Such biochemical adaptation is also accompanied, in toothed cetaceans, by morphological changes: such as enlarged mitochondria and midpiece (Alves et al 2020).

### Duplication and Loss: a “*domino effect*” links disruption of *Atp1a4* in separate mammalian lineages

We describe a striking case of parallel *Atp1a4* gene erosion, comprising species from Odontoceti cetaceans, pangolins and the naked mole-rat. Notably, most of these species were showed inactivating mutations in genes associated with sperm-specific glycolysis and encoding key routes of glucose subsidiary pathways (Figure 4; Alves et al. 2021). Why and how were these two evolutionary events linked? In mouse, deletion of the *Atp1a4* reduces sperm motility and hyperactivation, linked to an abnormal flagellar shape, leading to severe infertility (Jimenez et al. 2011; McDermott et al. 2021). Furthermore, *Atp1a4*-deficient sperm cells also displayed an accumulation of intracellular Na^+^, a depolarized plasma membrane, and a reduced capacity to regulate osmotic balance, partially underlying the observed morphological defects (Jimenez et al. 2011; McDermott et al. 2021). In this scenario, the male sterility observed in the mouse knock-out appears paradoxical in light of the findings reported here, as it suggests the essentiality of *Atp1a4* in mammals (Syeda et al. 2020). Conversely, two alternative scenarios can be envisioned to accommodate the adaptive loss of *Atp1a4*. A first setting would favour the maintenance of *Atp1a4* function, which would be co-opted to an alternative *Atp1a* gene paralogue: such as the *Atp1a1*, which displays a residual activity in sperm cells (Wagoner et al. 2005). In this case, the function would be maintained but no longer performed by the same Na^+^/K^+^-ATPase. Yet, in this scenario, we would expect the stochastic lineage loss of one of the two Na^+^/K^+^-ATPase, instead of the targeted erosion of the specific testis paralogue, *Atp1a4*. An alternative scenario includes the erosion of *Atp1a4* with a significant decrease of glucose uptake, implying the rewiring of sperm cell physiology in these lineages, notably glucose metabolism. In effect, Numata et al. (2022a; 2022b) have recently highlighted the role of a sodium dependent glucose transporter (SGLT1) in sperm function and motility. Furthermore, they showed that *Atp1a4*-deficient mice exhibited a decrease in glucose uptake and intracellular ATP: establishing a link between glucose transport and ATP1A4 activity (Numata et al. 2022b). In this context, the suggested loss of the testis glycolytic pathway in some mammal species (Alves et al. 2021), likely “*targeted*” the disruption of this specific Na^+^/K^+^-ATPase - mirroring a molecular “*domino*” (Dagan et al. 2006). This second hypothesis provides a more plausible explanation, since the erosion of *Atp1a4* in the individual lineages does not appear random but rather correlated with the suggested loss of sperm specific glycolysis. What would be the consequences of a complete *Atp1a4* deletion without an auxiliary mechanism? Although the deletion of the *Atp1a4* gene results in complete infertility of male mice, due to a severe reduction in total and hyperactive sperm motility (Jimenez et al. 2010; Syeda et al. 2020), the viability of *Atp1a4*-deficient sperm was significantly improved in hypertonic conditions, signifying that the external medium could abrogate some of the deleterious effects of the gene removal (McDermott et al. 2021). Thus, we hypothesize that *Atp1a4* functional loss in toothed-whales, pangolins and the naked mole rat was most likely a consequence of physiological changes, notably in sperm energy metabolism, along with the maintenance of the ubiquitous isoform of Na^+^/K^+^-ATPase (Blanco and Mercer 1998). Overall, the loss of *Atp1a4* in these 3 mammal lineages provides a remarkable case study of parallel evolution, illustrating how gene loss shapes the co-evolution of associated pathways towards novel physiological equilibria.

**Figure 4.**
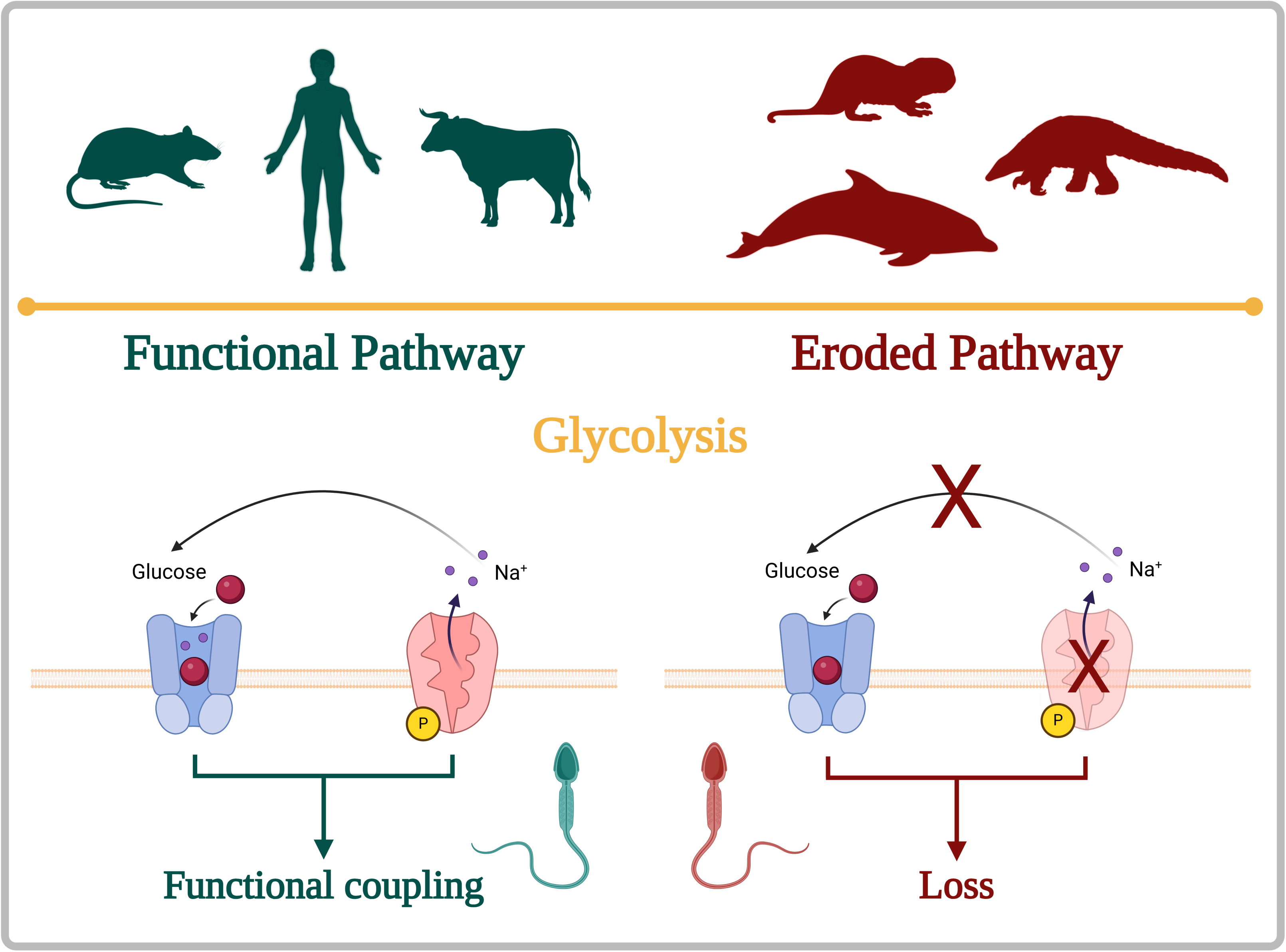
The evolution of glucose-based energy physiology in mammalian sperm cells. In mammal sperm cells, the loss of the glycolytic pathway reported in a few groups of species cancelled the functional coupling of a glucose transporter (SGLT1) the Na^+^/K^+^ ATPase (ATP1A4) responsible for maintaining an appropriate electrochemical balance between the intracellular and extracellular environments.

## Material and Methods

### Synteny and Phylogenetic analyses

Synteny and phylogenetic analyses were performed to inspect one-to-one orthology and clarify the timing of *Atp1a4* emergence. The synteny analysis was performed using *Atp1a4* from humans (*Homo sapiens*), chickens (*Gallus gallus*), green turtles (*Chelonia mydas*) and platypusus (*Ornithorhynchus anatinus*) genomes (*H. sapiens* - GCF_000001405.40, *G. gallus* - GCF_016699485.2, *C. mydas* - GCF_015237465.2 and *O. anatinus* - GCF_004115215.2).

To explore the *Atp1a4* gene evolution in mammals, a phylogenetic analysis was conducted. Nucleotide coding sequences of *Atp1a2* and *Atp1a4* were obtained from the NCBI for a representative group of species from major mammalian lineages and a selection of bird and reptile species. The sequences were aligned by translation aligned Geneious Primer software (Geneious 2021.2.2). The alignment was manually inspected and partial sequences were removed, as well as columns with more than 90% gaps. The resulting alignment was submitted to the PhyML3.0 server (Guindon et al. 2010) using the smart model selection method to automatically infer the evolutionary model (Lefort et al. 2017). Branch support for each phylogenetic tree was determined using standard bootstrap analyses (with 1000 bootstrap replicates). The resulting trees were visualized and analysed using FigTree V1.4.4, available at http://tree.bio.ed.ac.uk/software/figtree/.

### Transcriptomic analysis

We conducted a global analyses of relative gene expression in platypus (*Ornithorhynchus anatinus*), employing the approach described previously by Páscoa et al. (2022). First, we downloaded the genome and annotations of *O. anatinus* (Accession number: GCF_004115215.2) and *M. domestica* (Accession number: GCF_027887165.1) from the NCBI genome browser. RNA-seq datasets from NCBI were collected and concatenated when multiple datasets were available for a specific tissue. Prior to conducting the relative gene expression analyses, we ensured the accuracy of the genomes and annotations using the (agat_convert_sp_gxf2gxf.pl) script and converted them from .gff to .gtf format with the (agat_convert_sp_gff2gtf.pl) script from the AGAT tool (Dainat et al. 2022). Subsequently, the RNA-seq datasets were mapped against each genome using the Hisat2 v.2.2.1 aligner (Kim et al. 2015; 2019), and relative gene expression was determined with the StringTie v.2.2.1 software (Pertea et al. 2015), following the authors’ protocol (Pertea et al. 2016). The resulting gene expression quantifications were represented in transcripts per million (TPM) and fragments per kilobase per million mapped fragments (FPKM). Finally, to assess the relative *Atp1a4* expression across multiple tissues in humans, mice, and bulls, we checked the VastDb database (Tapial et al. 2017).

### Sequence extraction

Genomic sequence retrieval for *Atp1a4* annotation in 202 mammal species was accomplished using two different approaches: a) for annotated genomes, including species with annotated *Atp1a4* and species where no target gene annotation was found, the Genomic Sequence Downloader (https://github.com/luisqalves/genomic_sequence_downloader.py) was used. The human (*Homo sapiens*) synteny for *Atp1a4* was used as input reference, and downstream and upstream flanking genes were only considered if their Gene type was “protein coding”. If the Genomic Sequence Downloader was unable to retrieve a genomic region that included the target gene, a manual search was carried out using reference genome assemblies present at NCBI. This was accomplished by using either the target gene or the flanking genes to determine the genomic region encompassing the physical location of the gene; b) for unannotated genomes, the genomic sequences were retrieved via blastn searches using as query both *B. taurus* (cattle) *Atp1a4* coding sequence (CDS) (or the *H. sapiens* ortholog for non-cetacean mammals) and the CDS’s of the flanking genes. The best genomic scaffold (presenting the highest query coverage and identity value) was collected from the resulting BLAST hits. When target genomes presented a low N50 contig, the first two genomic scaffolds retrieved via blastn were considered for subsequent analyses.

### Gene inactivation inference

For gene annotation, the collected genomic sequences were uploaded to Pseudo*Checker* (http://pseudochecker.ciimar.up.pt), an integrated computational pipeline for gene loss deduction (Alves et al. 2020). Pseudo*Checker* analyses ran using default parameters and *B. taurus Atp1a4* nucleotide sequence (NCBI Accession ID: NM_001144103.2) or the *H. sapiens* orthologue (NM_144699.4) as references for Cetacea and non-Cetacea *Atp1a4* annotations, respectively.

Estimation of the disruptive condition of the tested gene was obtained through PseudoIndex - a user assistant metric built into the Pseudo*Checker* pipeline - varying on a discrete scale from 0 (coding) to 5 (inactivated). For each species, whenever the PseudoIndex was higher than 2, the genomic sequence was further imported into Geneious Prime for manual annotation and validation (Lopes-Marques et al. 2017). Succinctly, we mapped the individual reference exons, by utilizing the built-in map to reference tool with the highest sensitivity parameter selected, onto the corresponding raw genomic nucleotide data of the studied species. For gene annotation, the same references orthologs were consistently used as in Pseudo*Checker*. The resulting aligned regions were screened for inactivating mutations including exon deletions, frameshift insertions and deletions, altered start and stop codons, premature stop codons and splice site mutations. Validation of open reading frame (ORF) disrupting mutations (one per species), was performed through blastn searches in two independent Sequence Read Archive (SRA) genomic projects (when available), using as query the exon exhibiting the selected mutation. For this, unassembled genomic sequencing reads were collected, uploaded to Geneious Prime and mapped to the corresponding exon, to determine whether the data from different individuals of the same species support the presence of the ORF abolishing mutation.

### Sample Collection, DNA Extraction, and PCR Amplification

For further mutational validation, we performed polymerase chain reaction (PCR) in a skin sample of the common bottlenose dolphin (*T. truncatus*) obtained through standard biopsy darting procedures (Mathews et al. 1988). Whole-genomic DNA was extracted according to the conventional phenol-chloroform protocol (Sambrook and Russel 2001) and transferred to −20 °C for long-term storage. To target exons 5 and 15 of *Atp1a4* gene, two specific primer pairs for PCR amplification were designed via Primer designing tool “Primer – BLAST” (NCBI). Cycling parameters were as follows: denaturation at 98 °C/10 s then 30 cycles of 98 °C/1 s, 64 °C/5 s, 72 °C/7 s and finally an elongation at 72 °C/1min. The amplified PCR products were run in a 2% agarose gel electrophoresis. The resulting bands of the expected size for each gene were cut and purified with GelPure (NzyTech), following the manufacturers’ protocol and sent for sequencing at Eurofins Genomics. The sequencing results were then analysed with Geneious Prime in order to confirm the amplified sequence.

### Animal sampling, RNA extraction, library construction, and sequencing for Transcriptomic Analysis

A testis sample was obtained after the death of a male striped dolphin stranded in the Mediterranean Sea, off the coast of Spain at GPS coordinates: 39°28′45″N, 0°19′23″W in 2023. After RNA extraction, ribosomal RNA was eliminated from the total RNA, succeeded by precipitation using ethanol. Following fragmentation, the first strand cDNA was synthesized from randomly primed hexamers. In the second strand cDNA synthesis, dUTPs were substituted with dTTPs in the reaction buffer. The directional library was prepared after undergoing processes such as end repair, A-tailing, adapter ligation, size selection, USER enzyme digestion, amplification, and purification. The library’s quality and quantity were assessed using Qubit and real-time PCR for quantification, and bioanalyzer for size distribution. Finally, the libraries underwent pair-read sequencing (150 bp) on an Illumina HiSeq 4000.

### Masking of repetitive elements, gene models predictions, genome annotation and transcriptomic analysis

Considering that there is currently no available annotation for the *S. coeruleoalba* genome (GenBank Assembly Accession: GCA_951394435.1), we adopted the pipeline outlined by Gomes-dos-Santos et al. (2023) to obtain this dataset. In short, we began by constructing a novel repeat library using RepeatModeler v.2.0.133 (Smit and Hubley 2015a), which was subsequently utilized to mask repetitive elements with RepeatMasker v.4.0.734 (Smit and Hubley 2015b). Following this, gene prediction was executed on the masked genome assembly through the implementation of the BRAKER2 pipeline v2.1.6 (Brůna et al. 2021).

This process involved the integration of RNA-Seq data and protein spliced alignments as extrinsic evidence. The RNA-Seq data was sourced from GenBank (NCBI SRA Accession ID: ERR11837479), subjected to quality trimming using Trimmomatic v.0.3839, and then aligned to the assembly using HISAT2 v.2.2.0. For the protein dataset, we incorporated proteomes from various origins, as listed. To refine and filter the BRAKER2 predictions, we employed AGAT v.0.8.0 (Dainat et al. 2020).

Functional annotation was carried out through two primary methods: InterProScan v.5.44.80 (Quevillon et al. 2005) and BLASTP searches against the RefSeq database (Pruitt et al. 2007). For homology searches, DIAMOND v.2.0.11.149 (Buchfink et al. 2015) was utilized. The quality assessment of the predicted proteins was conducted using BUSCO v.5.2.2 (Manni et al. 2021) scores. Reports for statistics of repetitive elements, structural and functional annotations are presented.

Finally, we follow the approach of Pinto et al. (2023) to retrieve testis transcriptomic evidence of the *Atp1a4* gene in *S. coeruleoalba*. Briefly, gene expression was calculated using the novel RNA-seq tuxedo protocol (Pertea et al. 2016) and represented in TPMs and FPKMs.

### Semen collection

Semen collection from two dolphins has been previously described (Sánchez-Calabuig et al. 2017). After collection, semen samples (while avoiding exposure to light) were diluted at a 1:1 ratio using a commercial extender, Beltsville Thawing Solution (BTS) (GVP, Zoitech Lab, Madrid, Spain), and frozen in a TRIS egg yolk-based buffer containing glycerol, resulting in a final glycerol concentration of 3% and a sperm concentration of 200 x 10= sperm/mL.Frozen-thawed semen samples from two different Asturian Valley bulls were provided by the Regional Service of Agrifood Research and Development (SERIDA), Gijón,

Spain.Four to five straws (0.25 mL) of each animal (same ejaculate) were thawed at 37 ℃ in a water bath for 40 s. Motile spermatozoa were selected by BoviPure™ gradient (Nidacon Laboratories AB, Göthenborg, Sweden) centrifuged for 10 min at 290 × g. The resulted pellet was then resuspended in Boviwash solution (Nidacon Laboratories AB, Göthenborg, Sweden) and centrifuged for 5 min at 290 × g. Pellets were frozen at -80°C or used to western blot analysis.

### Western blot analysis of *Atp1a4* in common bottlenose dolphin spermatozoa

Semen samples from both dolphins and bulls were obtained from frozen straws. The total sperm count was determined using a Neubauer chamber, and samples were standardized to 20 x 10^^6^ total sperm per sample. Protein extraction was performed using 40 µL of RIPA buffer containing 50 mM Tris-HCl (pH 7.6), 150 mM NaCl, 1% Triton X-100, 0.5% sodium deoxycholate, and 0.1% sodium dodecyl sulfate (SDS), supplemented with Complete™ EDTA-free Protease Inhibitor Cocktail (11873580001, Roche) for 1 hour at 4°C, with vortexing every 10 minutes. The lysate was then centrifuged at 12,000 × g for 10 minutes at 4°C, and the supernatant was collected for protein analysis. Total protein content was quantified using the Qubit Protein Assay Kit (Q33211, Thermo Fisher) following the manufacturer’s protocol. Thirty-five micrograms of total protein from each sample in RIPA buffer were mixed with 2 µL of 4X NuPAGE LDS loading buffer (NP0008, Thermo Fisher Scientific) and 2 µL of DTT (500 mM), achieving a final volume of 10 µL per sample. The mixture was boiled at 70°C for 10 minutes. Proteins were separated on a 12% SDS-PAGE gel, with a molecular weight marker (Rainbow High Range, RPN756E, Cytiva) loaded as a reference. The proteins were then transferred onto a nitrocellulose membrane using a semi-dry high-molecular-weight transfer protocol. The membrane was stained with Ponceau S Red to confirm to confirm transfer, sample loading consistency and protein presence. After washing with PBST (0.05% tween-20 phosphate buffered saline, pH 7.4) immunostaining was performed. The primary antibody used as loading control was mouse anti-α-Tubulin (926-42213, Li-COR Inc, NE, USA).

Blocking was conducted with 5 mL of commercial Western blot blocking buffer (Fish Gelatin, T7131A, Takara) for 45 minutes. The membrane was incubated overnight at 4°C with constant agitation in rabbit anti-ATP1A4 polyclonal antibody (PA5-109429, Invitrogen) diluted 1:1000 in blocking solution. This was followed by incubation with the secondary antibody, Goat Anti-Rabbit IgG (H+L) HRP conjugated (bs-0295G-HRP, Bios Antibody), diluted 1:5000 in PBST, for 1 hour at room temperature with agitation. Chemiluminescence detection was performed using the Immobilon Forte Western HRP Substrate (WBKLS0050, Millipore). The chemiluminescence signal was digitized using an ImageQuant LAS 500 chemiluminescence CCD camera (GE Healthcare Life Sciences, USA, 29005063).

### Relaxed Selection Analyses

A dataset of 135 *Atp1a4* mammalian sequences was obtained from the NCBI assembled genome database. To analyse the selection pressure acting on the *Atp1a4* sequences in a context of gene integrity/erosion, we used mammal species for which we were able to assemble a full-length sequence; for species presenting a predicted non-coding *Atp1a4*, premature stop codons and frameshift mutations were removed. All sequences are provided in fasta format. Coding and corrected non-coding *Atp1a4* sequences were uploaded into Geneious Prime and aligned using translation align option. To test whether selection underwent relaxation in putatively deficient *Atp1a4* orthologs, we used the RELAX tool (Wertheim et al. 2015) implemented in the HyPhy v.2.5.25 package (Pond et al. 2005) with default parameters. We examined two test groups, i.e., cetaceans (vs. other mammals) and naked mole-rat (vs. other mammals). In both tests, the branches including species with an intact target gene were used as background.

### RNA-seq enrichment analysis of naked mole-rat and striped dolphin

We employed a comparative transcriptomic approach to explore the key differences between the testis of naked mole-rat (*H. glaber*) and striped dolphin (*S. coeruleoalba*) versus mouse (*M. musculus*) and bull (*B. taurus*) (Pinto et al. 2023). Briefly, a reference-based approach was employed to measure relative gene expression in *H. glaber* and *M. musculus*. Both genome assemblies and annotation files for each species were obtained from the NCBI database (https://www.ncbi.nlm.nih.gov/genome/). Transcriptomic datasets from testes were retrieved from the Sequence Read Archive (SRA) database (NCBI Accession Numbers; (*H. glaber* - SRR2120770; *M. musculus* - SRR5047954; *B. taurus* - SRR11448216)). Gene expression was measured using the same approach as described above in the *S. coeruleoalba* testis transcriptome, and for *H. glaber*, *M. musculus* and *B. taurus*, only genes with more than ten transcripts per million (TPM) were selected for further analyses.

To ensure data uniformity, the gene accessions with considerable gene expression (TPM>10) for each species were uploaded to g:Orth tool from the g:Profiler website (Raudvere et al. 2019; https://biit.cs.ut.ee/gprofiler/gost) to provide a list of *Homo sapiens* gene orthologs for each species. These gene lists were then subjected to enrichment analyses using the g:Profiler. Specifically, we performed the analyses for Gene Ontology (GO) molecular function (MF), GO cellular component (CC), and GO biological process (BP) with a statistical significance threshold of <0.05. To gain biological insights, we divided the results into three sub-tables: GOs enriched in both *H. glaber*/*S. coeruleoalba* and *M. musculus*/*B. taurus*, GOs exclusively enriched in *H. glaber*/*S. coeruleoalba*, and GOs exclusively enriched in *M. musculus*/*B. taurus*. The gene content of the top 10 enriched GOs, per category (MF, CC, BP), was analysed, particularly focusing on those uniquely enriched in *M.* musculus/*B. taurus*. Results from enrichment analysis between *H. glaber* vs *M. musculus* and *S. coeruleoalba* vs *B. taurus* can be found.

## Data and Resource Availability

The data that support the findings of this study are openly available in the National Center for Biotechnology Information (NCBI) Assembly database (https://www.ncbi.nlm.nih.gov/assembly/), as well as NCBI Sequence Read Archive (SRA) (https://www.ncbi.nlm.nih.gov/sra/). Genome assemblies are also available at DNA Zoo (https://www.dnazoo.org/) and at The Bowhead Whale Genome Resource (http://www.bowhead-whale.org/). The specific accession numbers (i.e., genome assemblies, scaffold IDs and SRA and sample IDs) for all data used from these publicly available databases.

The raw read sequencing outputs from *S. coeruleoalba* testis transcriptome were deposited at the NCBI Sequence Read Archive with the accession’s numbers: SRR30991393-SRR30991394. *S. coeruleoalba* repeat masked genome assembly and BRAKER2 prediction statistic are available. *S. coeruleoalba* genome annotation, prediction .gff files, as well as all predicted genes, transcripts and amino acid sequence files are available at Figshare: 10.6084/m9.figshare.28254077.

## Acknowledgments

We acknowledge the various genome consortiums for sequencing and assembling the genomes.

